# Tracking savanna vegetation structure in South Africa by extension of GEDI canopy metrics with Landsat, Sentinel-2, and PALSAR

**DOI:** 10.1101/2025.07.16.664478

**Authors:** Steven K. Filippelli, Jody C. Vogeler, Francisco Mauro, Corli Coetsee, Patrick A. Fekety, Melissa McHale, David Bunn

## Abstract

Adaptive management of savanna ecosystems requires frequent monitoring of woody vegetation structure, and although vegetation structure and changes may be captured with repeat airborne lidar, it is spatially and temporally limited across African savannas. As an alternative, this study evaluates the extension of spaceborne waveform lidar canopy metrics (RH98, Cover, Foliage Height Diversity) from the Global Ecosystem Dynamics Investigation (GEDI) across the Greater Kruger region in South Africa using moderate resolution optical sensors (Landsat and Harmonized Landsat Sentinel-2 [HLS]), L-band Synthetic Aperture Radar (PALSAR-1 and -2), and topographic and soil covariates. We compared the performance of 14 predictor sets incorporating different sensor combinations and temporal processing methods (LandTrendr and CCDC) in random forest models using temporal cross-validation to assess extrapolation accuracy. The most parsimonious fusion model (LandTrendr + SAR + topography/soils) achieved RMSEs of 3.04 m for RH98, 13.38% for Cover and 0.34 for FHD, which was comparable to more complex models using HLS and CCDC. All models demonstrated good temporal transferability with minimal bias but tended to overestimate low values and underestimate high values, which muted the estimated magnitude of change. Annual canopy structure maps derived from the best model captured expected spatial patterns and were used in model-based estimators to quantify changes in areas impacted by elephants, timber harvesting, fuelwood extraction, and woody encroachment. Extending GEDI metrics with moderate-resolution sensors thus offers a viable approach for large-scale savanna monitoring and detecting change in high impact areas.

## 2 ntroduction

Understanding the structure and dynamics of woody vegetation is crucial for ecological management and conservation of savanna biomes, which cover approximately a fifth of the Earth’s land surface and provide ecosystem services to over one billion people (Scholes and Archer, 1997; Twine, 2019). Frequent monitoring of woody structure is particularly important in savannas due to their highly dynamic nature, one that is strongly influenced by disturbance regimes. Significantly, these changes can have cascading effects on biodiversity (McCleery et al., 2018) and ecosystem function and services (Ding and Eldridge, 2024). In African savannas, remote sensing has become an indispensable tool for mapping and monitoring woody vegetation structure and its changes over time, with studies demonstrating its effectiveness in quantifying tree cover, canopy height, and biomass at various scales (see review by Abdi et al., 2022). While airborne lidar has been regarded as the most accurate remote sensing system for mapping canopy structure (Coops et al., 2021), its lack of spatial and temporal availability, particularly in African savannas, limit its application as a regular monitoring tool. Spaceborne lidar systems, like the Global Ecosystem Dynamics Investigation (GEDI), now enable regular, large-scale monitoring of woody structure across Africa’s vast and often inaccessible savannas (Abdi et al., 2022; Li et al., 2023). This advancement provides critical data for addressing key conservation challenges such as woody encroachment, megafauna impacts, habitat fragmentation, and resource extraction pressures.

The region of South Africa encompassing Kruger National Park (KNP) and adjacent private nature reserves has been a focal point for studying the natural and anthropogenic factors that shape vegetation structure in savannas. Changes in elephant populations, rewilding efforts, fuelwood and timber harvesting, woody encroachment, and illegal mining are only a subset of the changes occurring in this landscape that are of interest to scientists, land managers, and policy makers alike. In KNP, elephant numbers have surged from about ∼9,000 in 1994 when culling ceased to ∼31,000 in 2020 (Ferreira et al., 2024), significantly accelerating tree fall, which can be six times higher in some landscapes open to elephants (Asner and Levick, 2012). After 1994, policy changes were introduced to encourage wider area rewilding efforts. These included the closing of artificial waterholes and fence removal between the park and private reserves, leading to rapid redistributions of elephant impacts (de Boer et al., 2015; Smit et al., 2007). Human activities also contribute to structural changes — fuelwood harvesting in communal areas may promote coppice growth (Shackleton et al., 2022; Twine and Holdo, 2016) and reduce the abundance of large stems and preferred species depending on harvest regimes (Matsika et al., 2013). Forest plantations in the western portion of the Greater Kruger area provide high socioeconomic value for the region but also reduce streamflow within a catchment zone where maintaining water production and quality is a high priority (Dye and Versfeld, 2007; GKSDP, 2020). Additionally, widespread woody encroachment, likely linked to rising CO_2_ levels or possibly regional changes in fire regimes or livestock management, has been observed as doubling in woody cover across land uses between 1940 and 2010 (Stevens et al., 2016). These dynamics underscore the complexity of woody structure change in this savanna landscape, further complicated by the heterogeneity and seasonal variability inherent to these ecosystems.

Quantifying the magnitude and spatial patterns of woody structure change associated with these different disturbance types and change drivers is crucial for adaptive management in the Greater Kruger region. To implement adaptive management strategies that maintain desired ecosystem states and biodiversity, managers need to know how factors such as elephants, fuelwood collection, timber harvesting, and woody encroachment are impacting woody vegetation in near-real time and across sensitive areas (Gaylard and Ferreira, 2011). Obtaining sufficient field data to accurately quantify such changes in small areas of interest using traditional estimation methods can be challenging and resource-intensive (Rao and Molina, 2015). Remote sensing-based maps provide a promising alternative for enabling comprehensive assessments of vegetation structure and its changes across spatial and temporal scales in savannas (Abdi et al., 2022).

Remote sensing-based maps of woody structure can also be leveraged to assess average levels of structure or change for specific areas of interest, often with lower uncertainty than assessments relying on field data alone (Ståhl et al., 2016). This is especially important in savanna ecosystems, which typically feature low to moderate tree cover and may have subtle changes compared to clear cutting in temperate forests. However, these factors also pose a challenge for remote sensing in savannas. The prediction error and potential biases of woody structure maps or their change may be comparable to or even exceed the magnitude of actual change (Reinhardt et al., 2020), and these issues may propagate when assessing average change for areas of interest (Filippelli et al., 2024). Furthermore, applying remote sensing models outside the temporal range of their training data, known as “temporal transferability”, introduces the potential for additional bias, which can lead to inaccurate conclusions about the amount or even the direction of change (Fekety et al., 2015; Filippelli et al., 2024). For these reasons, it is imperative to check for model bias and quantify the uncertainty associated with remote sensing-based estimates of canopy structure and change to ensure that observed changes are real and not artifacts of model error or bias.

Previous research on remote sensing of woody structure in African savannas has provided valuable insights into methods for improving the precision of structure maps that can elucidate spatial patterns and ecosystem dynamics. Airborne lidar mapping of experimental herbivore exclosures and historical burn plots in KNP have revealed how geologic substrate, soil nutrient availability and moisture, fire regimes, and herbivory interact to influence canopy height and cover (Asner et al., 2009; Smit et al., 2010) and tree spatial patterns in savannas (Staver et al., 2019). Repeat lidar can provide greater detail and mechanistic understanding of these changes, for example by quantifying elephant treefall rates (Asner and Levick, 2012). Although limited in spatial and temporal availability, airborne lidar has also served to calibrate models based on moderate resolution sensors for broader scale monitoring. L-band SAR from the dry season has proven to be the most effective sensor for mapping of woody cover and biomass, with modest improvement in accuracy provided by fusion with optical sensors or C- and X-band SAR (Naidoo et al., 2016, 2015; Urbazaev et al., 2015). Changes in woody cover can be monitored through comparison of annual map products developed from L-band SAR, though use of direct backscatter change yields less underestimation of change and generally lower RMSE and bias (Wessels et al., 2023). These examples illustrate the critical role played by airborne lidar in helping to understand savannas at experimental sites or through extension with moderate resolution sensors. However, they are limited to the vegetation types and time periods captured in the airborne lidar. There is still a need for more frequent monitoring of woody structure over a greater diversity of vegetation types within savannas to enable adaptive management to rapidly respond to changes as they are occurring.

Sampling spaceborne lidar systems, such as GEDI, provide a potential alternative for frequent monitoring woody structure over a greater diversity of landscapes due to their spatially extensive multi-year coverage. In southern African savannas, GEDI canopy height captured during leaf-on had an RMSE of 1.64 m and bias of -0.55 m when compared to waveform simulation with airborne lidar (Li et al., 2023), proving its potential utility as an alternative reference source. GEDI relative height 95^th^ percentile (RH95) was extrapolated with Landsat to produce a 2019 global canopy height map at 30 m resolution with an overall model RMSE of 6.6 m, an airborne lidar validation RMSE of 9.07 m, and no airborne lidar validation in savannas (Potapov et al., 2021). GEDI L2A height metrics and L2B canopy metrics have also been expanded wall-to-wall with moderate resolution sensors in western US forests (Vogeler et al., 2023) and other regions (Geremew et al., 2024; Kacic et al., 2021), but applications in savanna systems remain limited. In one example, modeling GEDI RH95 with Landsat, Sentinel-1, and Sentinel-2 in western Australia eucalypt savannas had an RMSE of 2.68 m (Lutz et al., 2024). There remains a need to evaluate the wall-to-wall extension of GEDI canopy metrics in African savannas as has been done in other regions.

The goal of this study was to test different remote sensing data sources and approaches for mapping GEDI canopy structure metrics across the Greater Kruger region over time to enable monitoring of woody structure changes. To this end, we specifically sought to answer three fundamental questions:

1. How do moderate resolution data source and processing methods influence the accuracy in mapping GEDI canopy metrics wall-to-wall in an African savanna?
2. How do different modeling approaches affect accuracy when temporally transferring the model to time periods outside that of GEDI’s temporal availability?
3. Do maps of canopy structure capture known spatial patterns and magnitude of change in areas with known elephant damage, timber removal, fuelwood harvesting, and woody encroachment?

To answer these questions, we modeled the GEDI L2B canopy metrics of RH98, canopy cover, and foliage height diversity with multiple predictor sets reflecting trade-offs in temporal availability and processing complexity. These included Landsat and Harmonized Landsat Sentinel-2 processed with the LandTrendr and Continuous Change Detection and Classification (CCDC) algorithms, PALSAR-1 and -2, topographic indices derived from the Shuttle Radar Topography Mission (SRTM), and predicted soils layers from the Innovative Solutions for Decision Agriculture Ltd. (iSDA). Maps of canopy structure for the Greater Kruger region were then used in model-based estimators to quantify changes in three areas of interest with known change drivers. The results of this study will form the basis for regular monitoring of woody structure in the Greater Kruger region and inform future research for spaceborne lidar supported vegetation monitoring in savannas.

## 3 Methods

### 3.1 Greater Kruger

South Africa’s Greater Kruger is a 60,600 km^2^ region defined in this study as encompassing KNP, a collection of private nature reserves and communal lands immediately to the west, and agricultural and urban development extending to the Drakensberg mountains in the west and the southern border of Ehlanzeni district municipality to the south. The geologic parent material, precipitation patterns, and topography are the primary drivers of vegetation patterns in this region. KNP has a distinctive divide of nutrient poor granitic soils in the western half of the park and nutrient rich basaltic soils in the east. Average annual precipitation follows a gradient of generally becoming wetter (750 mm/yr) when moving towards the north or west and drier (425 mm/yr) in the south and east. The savannas in the eastern portion of Greater Kruger exist on low undulating hills that lead to flatter grassy plains going further east (150 m to 750 m in KNP), while greater relief is present in the west rising to forested mountains (up to 2000 m).

The Lowveld ecozone, which covers KNP and the immediately surrounding area, contains 35 vegetation types (Gertenbach, 1983). The most prevalent woody vegetation found across these classes include *Colophospermum mopane* dominating vast areas of the north, *Combretum spp.* more often dominating in the south, and *Terminalia spp.* covering microtopographic patches (Figure 1). Vegetation characteristics within the eastern granitic landscapes follow a topographic gradient known as a catena with different woody species typically found along sandy crests, wet seep lines, midslopes, and footslopes. Although Lowveld savannas are dominated by shrubs and short stature trees of ∼2-5 m, there are sparsely interspersed tall trees (>5 m) that emerge from this matrix (e.g., *Sclerocarya birrea*, *Senegalia nigrescens*, *Adansonia digitata*). Weaving through the savanna are also riparian forests consisting of taller trees with broad crowns (e.g., *Diospyros mespiliformis*, *Ficus sycomorus*).

**Figure 1.**
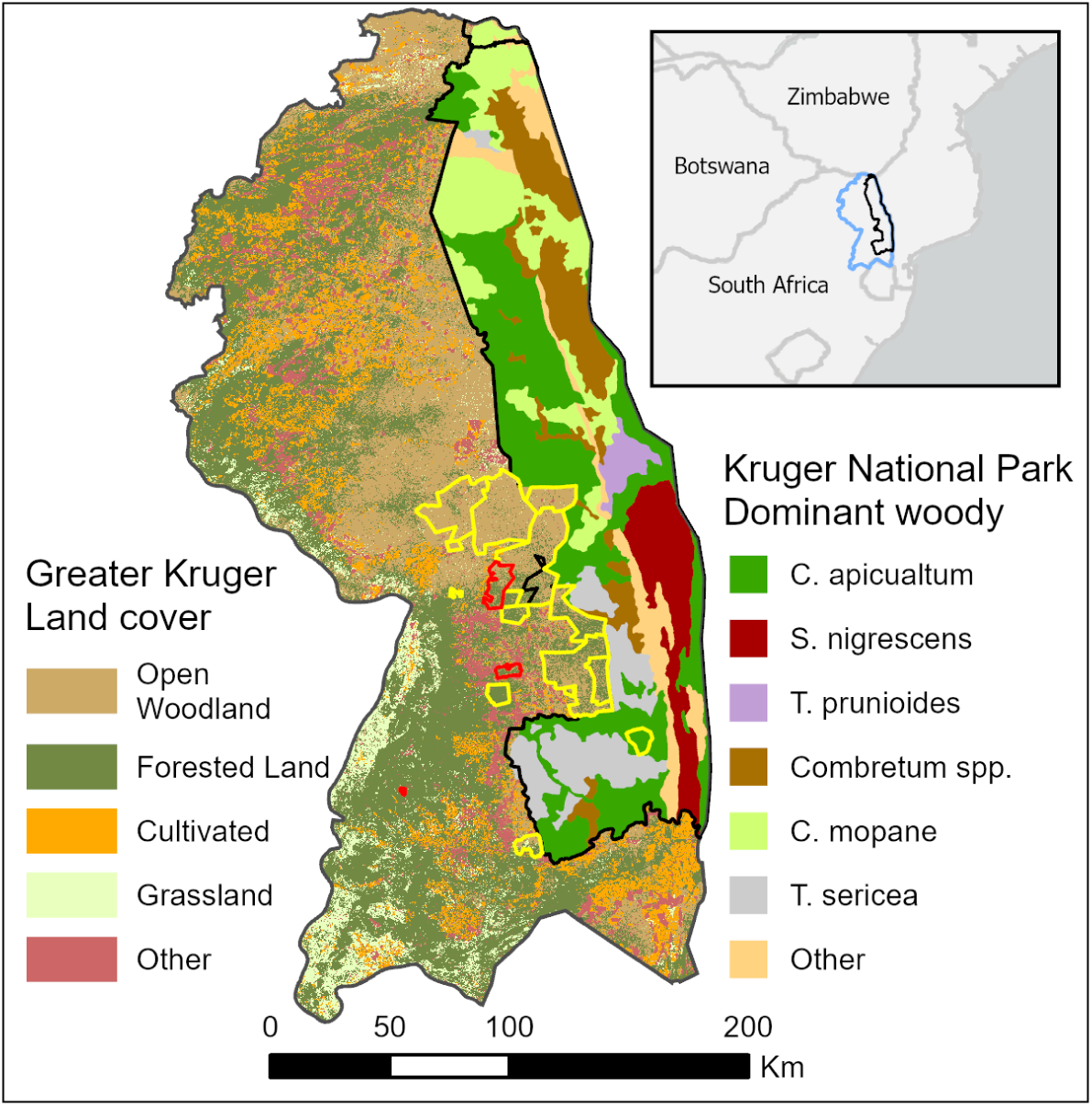
The study area as defined by the Greater Kruger Strategic Development Programme is shown with the 2020 South African National Land Cover Level 1 classification but including the more detailed Open Woodland class which also covers most of Kruger National Park. For Kruger National Park in the eastern half of the study area, the diversity of vegetation types is portrayed using the dominant woody vegetation species from a simplification of the Venter 1990 land systems. Areas used in small area estimation are outlined in yellow and the subset of case study areas are outlined in red.

### 3.2 Mapping GEDI canopy metrics

#### 3.2.1 GEDI data

We acquired the GEDI Level 2A Geolocated Elevation and Height Metrics version 2 product (GEDI02_A) and the GEDI Level 2B Canopy Cover and Vertical Profile Metrics version 2 product (GEDI02_B) for all shots collected over the study area between January 1, 2019 and December 31, 2023. These data were filtered to retain shots that had an L2A and L2B quality flag of 1 and that were collected during leaf-on (121 to 305 days of year). Li et al. (2023) found phenology to have the largest impact on GEDI canopy height accuracy in African savannas, whereas the use of coverage versus power beams or day versus night shots made no difference so long as the sensitivity was at least 90%. We extracted the relative height 98th percentile (RH98) metric from the L2A product and canopy cover (Cover) and foliage height diversity (FHD) from the L2B product. RH98 is the 98^th^ percentile relative height above the perceived ground elevation and is the most commonly used metric of canopy height to reduce the influence of outliers. The GEDI Cover metric can be interpreted as “the percent of the ground covered by the vertical projection of canopy material” (Tang and Armston, 2019). FHD is a equivalent to the Shannon diversity index for vertical heterogeneity of canopy elements, which is calculated by taking the proportion of leaf area index in a vertical layer and multiplying by its log and then taking the negative of the sum across all vertical layers (Tang and Armston, 2019). In African savannas, GEDI RH98 collected during leaf-on had an RMSE of 1.48 m and a bias of -0.55 m when compared to airborne lidar simulated waveforms (Li et al., 2023). The FHD and Cover metrics from GEDI have not been compared to alternative measurements of these metrics in savannas.

We further compared GEDI RH98 to field measurements of maximum canopy height at select locations across our study area. An opportunistic sample of 26 GEDI footprints in Kruger National Park were visited in January 2023, May 2023, and May 2024. We navigated to the center of the footprint with a Trimble GeoXH, which has a rated precision of 10 cm, and then used a laser range finder to measure the maximum canopy height within the footprint area.

**Figure 2.**
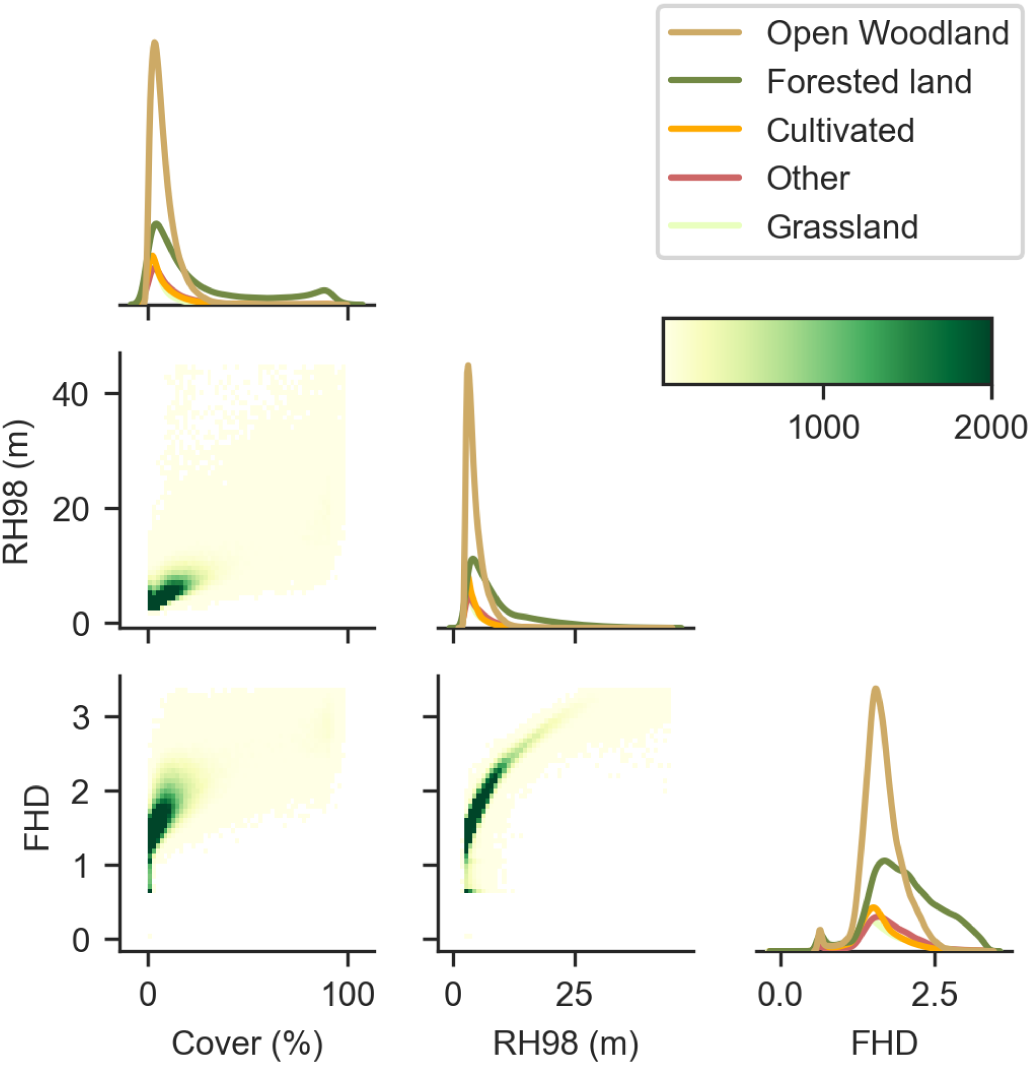
A pair plot showing the distributions of the GEDI canopy metrics used modeling and their relationship to each other. Kernel density estimates of the individual metrics are divided by the intersecting 2020 South African National Land Cover Level 1 classification. The pairwise relationships between metrics are shown by 2-D histograms.

#### 3.2.2 Spatial predictors

We evaluated Landsat, Harmonized Landsat Sentinel-2 (HLS) and PALSAR-1/2 data as remote sensing predictors in models of the GEDI canopy metrics. The Landsat Collection 2 Level 2 Tier 1 surface reflectance product for 1984-2023 and the HLS 30 m surface reflectance data for 2013-2023 were obtained and processed in Google Earth Engine (GEE) (Gorelick et al., 2017). Because only the Landsat portion (HLS L30) of the HLS product was available in GEE, we created the Sentinel-2 portion of the dataset by applying the same coefficients for bandpass adjustment used by the HLS product and used area-weighted mean resampling to 30 m. We did not apply the Nadir Bidirectional Reflectance Distribution Function (NBAR) normalization to the Sentinel-2 data, and the cloud probability layer created with the sentinel2-cloud-detector was used for cloud masking with a threshold of 0.65. The Landsat scenes were masked to exclude clouds, cloud shadows, aerosols, and radiometric saturation using flags in the pixel quality assurance band created through the Fmask algorithm. We calculated several spectral indices for each dataset including the Normalized Difference Vegetation Index (NDVI) (Rouse et al., 1974), the Normalized Burn Ratio (NBR) (Key and Benson, 2006), the Normalized Difference Moisture Index (NDMI) (Gao, 1996), Tasseled Cap Brightness (TCB), Tasseled Cap Greenness (TCG), and Tasseled Cap Wetness (TCW) (Crist, 1985; Huang et al., 2002).

The Landsat and HLS data were then processed with two time-series algorithms for comparison: the Continuous Change Detection and Classification (CCDC) algorithm (Zhu and Woodcock, 2014) and the LandTrendr algorithm (Kennedy et al., 2018, 2010). CCDC generates harmonic regression coefficients for each pixel and band while automatically creating break points to account for disturbances. Coefficients from CCDC capture the overall brightness and phenology of observed surface reflectance as well as its trend over time. We extracted these coefficients from the end of the year (December 31) to represent the conditions of that year. We also created synthetic reflectance values representing at the middle of the wet season (February 1^st^) and dry season (September 15^th^) to account for changes in canopy structure that occur between years but within a CCDC segment that spans multiple years. For running CCDC, we excluded scenes with greater than 80% cloud cover to reduce the possible influence of unmasked clouds. Previous large area monitoring projects have applied CCDC to only the original spectral bands (Brown et al., 2020), and so we excluded spectral indices from CCDC outputs to reduce the number of variables and processing time. LandTrendr applies a linear fit to annual composites and identifies breakpoints to create linear temporal segments that reduce between year variations that are likely not due to actual changes in canopy structure (Kennedy et al., 2010). For processing each band and spectral index with LandTrendr, we first applied temporal compositing by calculating the medoid for the wet season (October 1^st^ to April 30^th^) and dry season (May 1^st^ to September 30th). Because LandTrendr is more susceptible to the influence of clouds, particularly at the beginning and end of the time series (Kennedy et al., 2010), we excluded scenes which had greater than 50% cloud cover. LandTrendr was run on each band and spectral index independently to produce “fitted” values using default parameters but with a maximum of 6 segments and prevent one year recover turned on.

The Global PALSAR-2/PALSAR version 2 annual mosaics of L-band SAR backscatter for 2007-2010 and 2015-2022 were also obtained from GEE. This 25 m resolution annual mosaic product is created from PALSAR strip maps collected in Fine Beam Dual mode and Wide swath Beam Dual mode with visual selection of strips to minimize the influence of surface moisture, ortho-rectification and slope correction with the ALOS World 3D digital surface model, and then radiometric balancing between adjacent strips. We converted these data from digital numbers to gamma naught and applied multitemporal speckle filtering using a boxcar spatial filter with a kernel size of 7 (Quegan and Yu, 2001), which was found to be effective in this region (Wessels et al., 2023). The HH and HV backscatter coefficients and local incidence angle were used as predictors of the canopy metrics.

In addition to the above temporally varying remote sensing predictors, we included temporally static spatial data on topography and soils as predictor variables in modeling the canopy metrics. A series of topographic metrics were created from elevation data collected by the Shuttle Radar Topography Mission (SRTM). Elevation, percent slope, aspect, the cosine of aspect, the sine of aspect, percent slope multiplied by the sine and cosine of aspect, transformed aspect, and topographic position index at 30 m, 300 m, and 990 m were all created from elevation data from NASADEM (NASA JPL, 2020), which is based primarily on SRTM. We also obtained the multiscale topographic position index, continuous heat-insolation load index, and topographic diversity index based on SRTM from Theobald et al. (2015). Predicted soils information at a 30 m resolution was obtained from the Innovative Solutions for Decision Agriculture Ltd. (iSDA) soils layers for mean organic carbon, cation exchange capacity, and nitrogen at 0-20 cm and 20-50 cm (Hengl et al., 2021).

#### 3.2.3 Modeling and mapping

For compiling model training and validation data, the area-weighted mean of each predictor layer was extracted for each GEDI footprint. For the temporally varying layers based on Landsat, HLS, and PALSAR, we extracted data for the year corresponding to the GEDI collection date. The wet season was associated with the year of its start date. For example, data for 2022 includes data obtained from the dry season dates of May 1, 2022 to September 30, 2022 and wet season dates October 1, 2022 to April 30, 2023. We spatially filtered GEDI shots to reduce spatial autocorrelation by randomly selecting at most one shot within 500 m square grid cells.

Twelve empirical models of the GEDI canopy metrics were developed from the above predictor data sets using random forest regression (Breiman, 2001). All random forest models used 100 trees and at each node the square root of the total number of variables was considered for splitting. The first two models used only Landsat predictor variables fitted with either LandTrendr or CCDC (CCDC^L30^) to enable hindcasting of vegetation structure for change analysis over longer time periods. The third model used HLS data with CCDC (CCDC^HLS^), which offers greater temporal frequency for filling temporal gaps in the cloudy wet season and reducing artificial breakpoints, but HLS data is limited to 2015 forward. A model was created with PALSAR data only, which is limited to 2007-2010 and 2015-forward, but provides a more direct measurement of woody biomass. We also tested the fusion of each of the optical data sources with PALSAR and the topographic and soil static variables.

Accuracy was assessed for all models by the root mean square error (RMSE), coefficient of determination (R^2^), and bias (i.e., mean of residuals) using out-of-bag predictions. We also implemented temporal cross-validation using withheld years of test data (Filippelli et al., 2024; Roberts et al., 2017) to estimate the accuracy of extending the models beyond the time period of the training data. Based on this accuracy assessment, we selected the best model for use in wall-to-wall annual mapping of the canopy metrics and subsequent analyses of change in woody structure.

### 3.3 Small area estimation for known areas of change

Maps of RH98, COV, and FHD were used to assess changes in canopy structure for areas of interest, which were primarily in private nature reserves along the western border of KNP. We delineated 18 areas of interest using a combination of property boundaries, very high-resolution satellite imagery, and descriptions in previous literature. Of these areas, we focus on four areas known to be affected by elephant damage, fuelwood extraction, timber harvesting, and woody encroachment. The first site, Thornybush Nature Reserve, removed its eastern fences in 2017 to create a continuous landscape with KNP, and the elephant population increased from 55 in 2017 to 770 in 2019 (Hannes Zowitsky, personal communication), which has resulted in high levels of tree damage and mortality. Second, an area of tree plantations that underwent a series of harvests starting in 2015 was delineated in the Drakensberg mountains to the west of Kruger National Park. Third, communal lands in the Bushbuckridge area have been subject to intense fuelwood extraction in some areas, and previous research has indicated that this has mostly resulted in coppice growth (Mograbi et al., 2015). The savanna regions of GKNP have in general been undergoing woody encroachment, which is believed to be caused by CO2 enrichment (Stevens et al., 2016). We delineated an area southeast of the Skukuza rest camp, which had noticeable woody encroachment between 2009 and 2024 Google Earth imagery.

Small area estimation was used to quantify the mean and construct confidence intervals of the canopy metrics for the areas of interest before and after the year the impacts occurred, or at the start and end of the canopy metrics map time series (i.e., 2007 and 2021). We used the Battese-Harter-Fuller unit-level model-based estimator (Battese et al., 1988; Rao and Molina, 2015) with the predicted canopy metric maps serving as the auxiliary data available for each unit of the population (i.e., a pixel). The model for the estimator was fit to the sample of GEDI shots used in the remote sensing model described above. All eighteen areas were used in estimation to reduce the standard errors by allowing the estimator of each area to “borrow strength” from the neighboring areas, and we combined areas with the years in which they contained GEDI footprints to form 62 domains that were used in model-based estimation. The Battese-Harter-Fuller model is based on a mixed effects model that uses independent areas as domains, and our models, which have temporal dependence, may not be as efficient in obtaining the precision of estimators. Because of this we also compared mixed effects models using area and time combined (i.e., the domains described above) with models using alternative random effects structures of area and area crossed with time.

We present small area estimation results on only four of the impact areas at two time periods for brevity and to illustrate how the canopy metric maps and model-based estimation could be used to quantify canopy changes. In years without GEDI (i.e., prior to 2018), the estimator of the mean is fully synthetic and thus more likely subject to bias. The temporal cross-validation described in the previous section is intended as a check for potential bias in the estimators when they rely on canopy maps created from applying the remote sensing model outside the temporal domain of the training data. The estimators for the mean for each domain were obtained with the sae package in R (Molina and Marhuenda, 2015). We constructed 90% confidence intervals from the mean squared error obtained from parametric bootstrapping with 200 bootstrap replicates (González-Manteiga et al., 2008), which was also obtained with the pbmseBHF function from the sae package (Molina and Marhuenda, 2015).

The model-based estimators were also evaluated for potential bias by comparison to the unbiased design-based direct estimators where sufficient GEDI footprints overlapped each area of interest in a given year (e.g., Thornybush reserve in 2019), which we consider as a single domain. The GEDI sampling design is effectively a one-stage cluster sample where each track is a cluster (Patterson et al., 2019). We obtained the direct estimator for the population mean of each domain with the survey package in R (Lumley, 2024) using a one-stage cluster design assuming a simple random sampling without replacement of clusters of unequal size. The 90% confidence intervals were also obtained with the survey package when at least two GEDI tracks intersected the domain to enable estimation of the variance.

## 4 Results

### 4.1 Models of GEDI canopy metrics

Random forest models of the GEDI canopy metrics had similar accuracy statistics between predictor sets using the same combination of sensor types despite using more complex processing (i.e., CCDC versus LandTrendr) or more observations (i.e., HLS versus L30) (Figure 3). For example, models using optical data sources alone (LandTrendr, CCDC^L30^, CCDC^HLS^) had an RMSE that differed at most 2.6% from the mean RMSE across the set of models for the same metric. The model RMSEs were nearly the same for CCDC^HLS^ than for CCDC^L30^, as were the CCDC fit RMSEs for each band with mean differences of 1.28e-3 for blue, 8.84e-4 for green, 2.47e-4 for red, 7.74e-4 for NIR, 1.34e-3 for SWIR1, and 1.37e-3 for SWIR2. Combining predictors from different types of data sources yielded greater improvement in models. For example, supplementing optical predictors with PALSAR reduced RMSE an average of 4.7%, and adding both PALSAR and soils and topography predictors reduced RMSE an average of 7.0%. Models using only optical predictors had higher performance than the SAR-only models. For example, the LandTrendr model explained 11.8%, 14.8%, and 8.7% more variance in cover, RH98, and FHD, respectively, than the PALSAR model.

**Figure 3.**
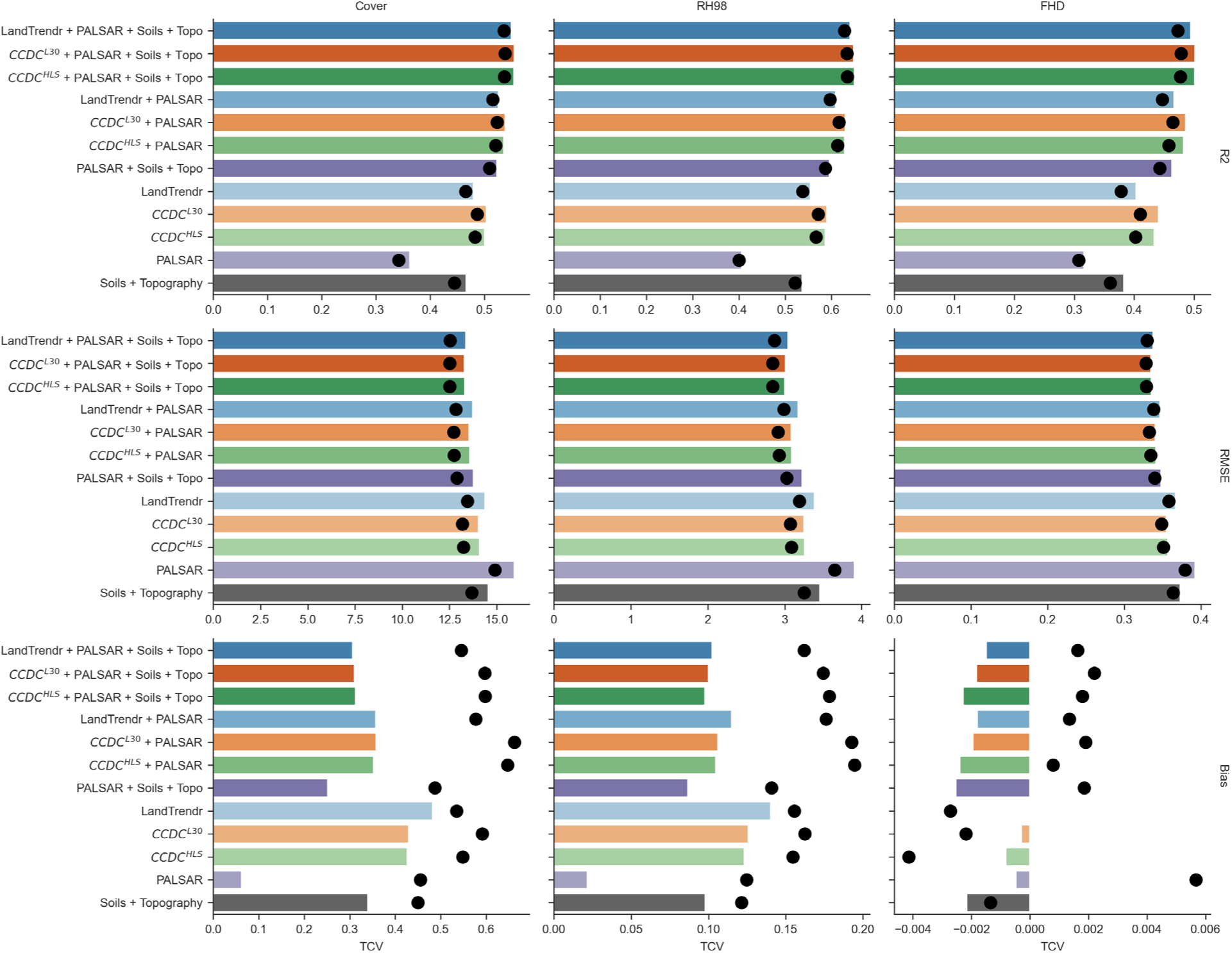
Bar plots showing the random forest out-of-bag model accuracy statistics of R2, RMSE and bias for each of the GEDI canopy metrics of Cover, RH98, and FHD for each the predictor sets. Black dots show the corresponding temporal cross-validation statistic.

Leave-one-year-out temporal cross-validation statistics (TCV) were very similar to the out-of-bag model statistics (OOB) using data from all years (Figure 3). TCV R^2^ was lower than the corresponding OOB R^2^ by 0.02 on average, and TCV RMSE was lower than OOB RMSE by 4.8% on average. While TCV bias appears to be significantly higher than OOB bias (Figure 3), both TCV bias and OOB bias were still close to zero across all models. Bias as a percentage of the mean observed value was at most 3.3% for the OOB models (LandTrendr Cover model) and 4.6% for TCV models (CCDC^L30^ + PALSAR Cover model).

The models using the LandTrendr + PALSAR + Soils + Topography predictor set (Figure 4) were chosen for mapping canopy structure across time because they had nearly the same accuracy as the most accurate models but were substantially less complex. These models using LandTrendr had RMSE values that were higher by only 0.07% for Cover, 0.04 m for RH98 and 0.003 for FHD in comparison to the most accurate models that instead used the CCDC^L30^ or CCDC^HLS^ predictor sets with 54 more variables. Although the models had near zero bias overall, there was substantial bias within particular ranges of observed values, reflecting the compression towards the mean that is common for random forest and other machine learning models (Zhang and Lu, 2012). The Cover model tended to overestimate the abundant low cover values, with an observed cover of <1% being predicted as 10.4% on average, and the rarer cover values over 50% were underestimated by 27.2% cover on average. This compression towards the mean was less extreme for RH98 and FHD.

**Figure 4.**
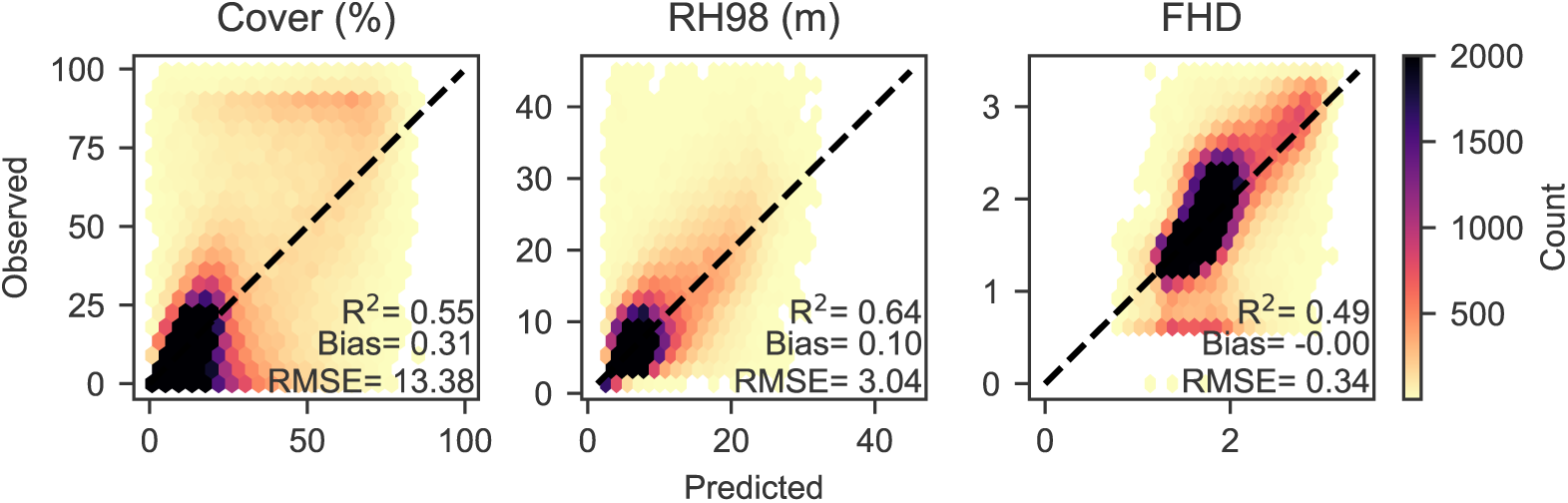
A hexagonal bin plot showing the two-dimensional distribution of observed and predicted values for each GEDI canopy metric along with the accuracy statistics for the models using the LandTrendr + PALSAR + Soils + Topography predictor set.

The compression towards the mean of predicted RH98 was also apparent when compared to field observations of maximum tree height within GEDI footprints (Figure 5). The regression of predicted RH98 on field maximum tree height had a slope of 1.94, but this was largely due to the leverage of a single tall tree. Without this observation, the linear regression became closer to 1:1 with a slope of 1.41 and intercept of -1.35, and the direct RMSE (i.e., not of the linear regression) dropped to 2.3 m. The GEDI footprint RH98 values corresponded closely with the field measured maximum tree heights, having a slope close to 1, intercept close to 0, and RMSE of 2 m.

**Figure 5.**
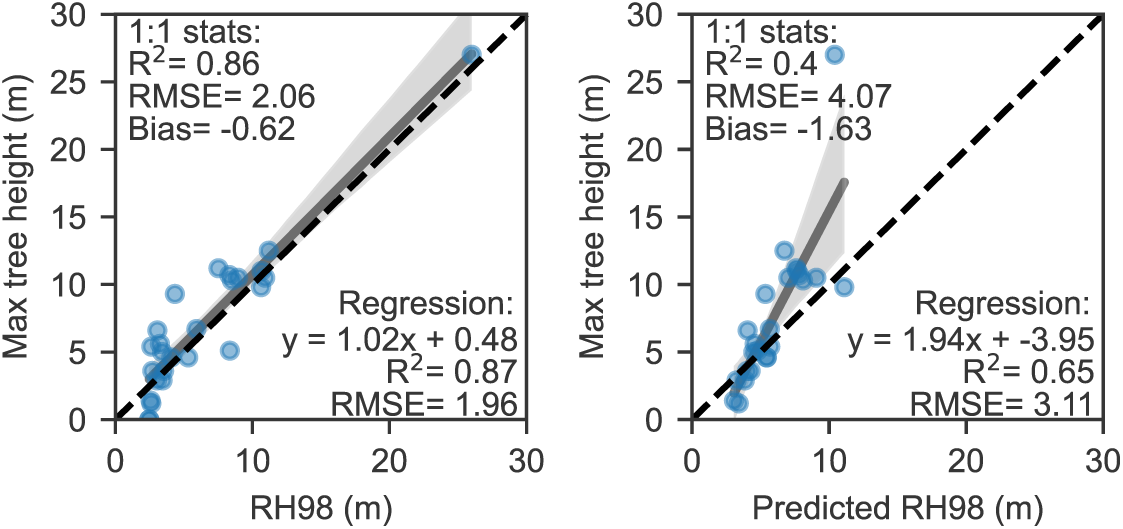
Comparison of field measured maximum tree height to GEDI footprint RH98 and predicted RH98 from the LandTrendr + PALSAR + Soils + Topography model. Statistics for this comparison are given for the linear regression of these values and with respect to the 1:1 line (i.e., direct comparison). Shaded grey area shows the 95% confidence interval.

The mixed effects models used in model-based estimation for each metric were fit to the sample of 15688 GEDI observations and random forest model predictions in the 62 area-year combinations (i.e., domains). Across domains there was a minimum of 2, maximum of 1413, and mean of 253 sample units. The fixed effects of these models were based on the random forest predicted value and often had coefficients close to 1: cover β = 0.94, RH98 β = 1.01, and FHD β = 1.00. The standard deviation of the random effects for the domains in were close to zero for RH98 (0.43) and FHD (0.048), which indicates there was little difference in the fit of predictions to observations between domains, but the random effect for the cover model had a relatively higher standard deviation of 2.3. In our comparison to models with alternative random effects structures, we found models using the combined Area:Time random effect had the lowest AIC (Table 1).

**Table 1.** Comparison of mixed effects models with different random effect structures using area, the cross of area and time, and area:time combined. The AIC and the standard deviation of the random effects are given for each model.

| Model | Area |  | Area + Time |  |  | Area:Time |  |
| --- | --- | --- | --- | --- | --- | --- | --- |
|  | AIC | Area | AIC | Area | Time | AIC | Area:Time |
| Cover | 106877 | 2.23 | 106716 | 2.14 | 1.00 | 106511 | 2.31 |
| RH98 | 64233 | 0.327 | 63995 | 0.320 | 0.318 | 63981 | 0.422 |
| FHD | 3700.6 | 0.0316 | 3598.1 | 0.0304 | 0.0320 | 3571.3 | 0.0473 |

For the 35 small area and year combinations with at least two sampling clusters (i.e., GEDI tracks), we compared the design-based estimators for the mean of each metric to the model-based estimators derived from the maps. The estimators mostly followed the 1:1 line, having R^2^’s of 0.8 to 0.88 (Figure 6a). The model-based estimator fell within the 90% confidence interval of the unbiased design-based estimator for 57%, 51%, and 49% of instances for cover, RH98, and FHD, respectively. The confidence intervals of both estimators overlapped in 77%, 80%, and 69% of instances for cover, RH98, and FHD, respectively. Despite the lack of overlap in confidence intervals, the average absolute percentage difference of the model-based estimator from the corresponding design-based estimator was only 8.7% for cover, 4.8% for RH98, and 1.8% for FHD. Although the differences between the estimators were typically small, Figure 6b shows that they mirrored the pattern of compression towards the mean similar to that of the model results in Figure 4. Domains with a design-based mean less than that of the mean of the training data (vertical grey dotted line in Figure 6b) tended to have a positive difference (i.e., possible overestimation by the model-based estimator), and those with a design-based mean greater than the training data mean tended to have a negative difference (i.e., possible underestimation by the model-based estimator). Really large differences between the estimators were in domains that only had two GEDI tracks (i.e., sample clusters), which could be part of the reason for the difference if the tracks sampled less representative portions of the domains.

**Figure 6.**
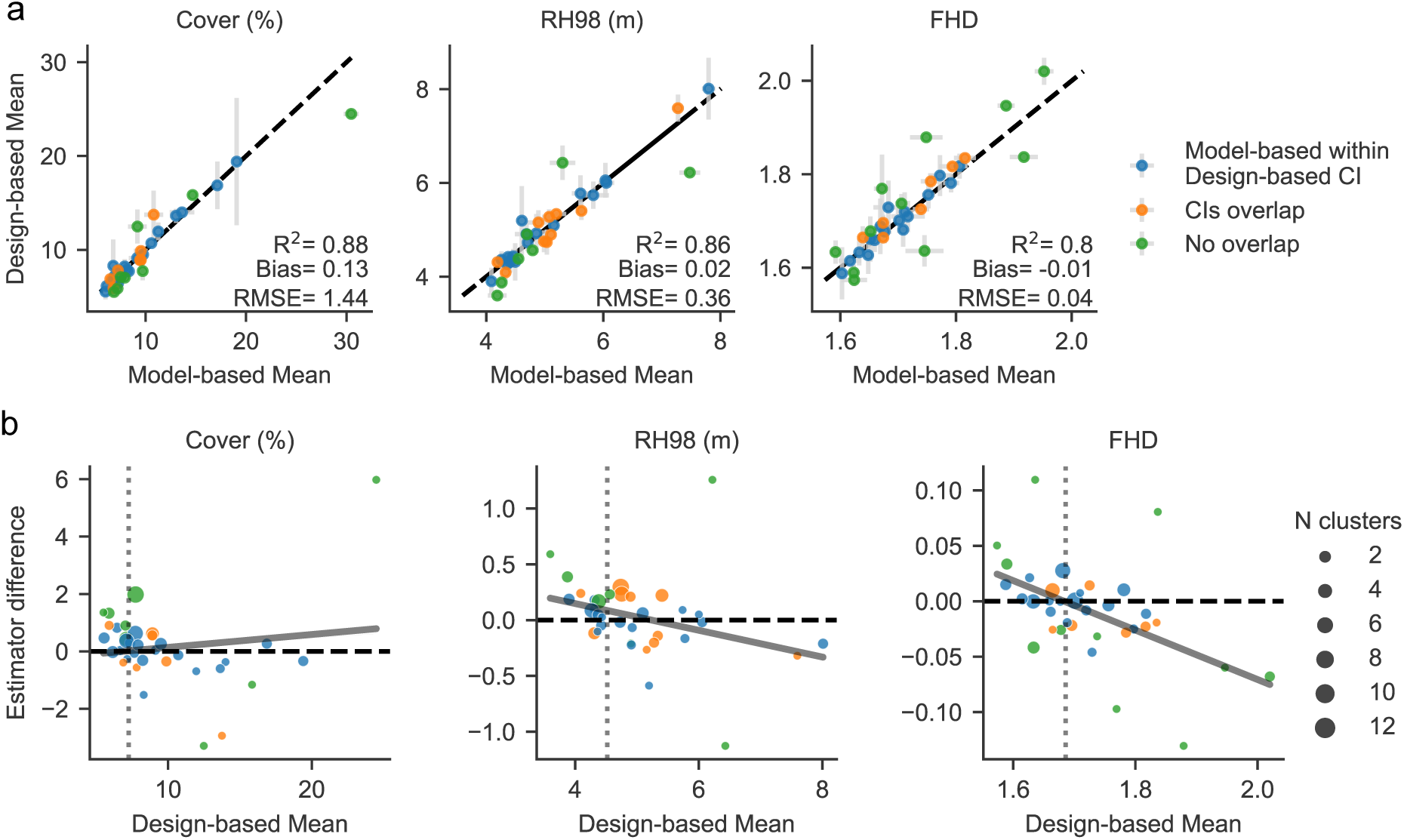
a) Comparison of model-based and design-based estimators for the mean of each GEDI metric and their 90% confidence intervals (grey bars). Accuracy statistics are given for the fit of the model-based estimators compared to the design-based estimators with respect to the 1:1 line (dashed black). Where the vertical grey bars cross the 1:1 line indicates where the model-based estimator falls within the confidence interval of the design-based estimator (blue dots), and where only the confidence intervals overlap are shown as orange dots. b) The difference of the model-based estimators from the design-based estimators are shown across the range of the design-based estimators to highlight the generally negative sloping pattern of the residuals. The vertical dotted grey line shows the mean value of the metric used in model training, which largely partitions the positive and negative residuals. The domains with large differences between the estimators were more likely to have only 2 sample clusters (i.e., GEDI tracks), which are shown as the size of each dot in (b).

### 4.2 Maps of canopy structure and change for impact areas

Maps of GEDI Cover, RH98, and FHD were created from models using the LandTrendr + PALSAR + Soils + Topography predictor set and used to quantify canopy structure changes in areas known to have undergone elephant impacts, timber harvesting, fuelwood harvesting, and woody encroachment between 2007 and 2022, when PALSAR-1 & 2 annual mosaics were available. The map of 2021 RH98 reflected the broader patterns of canopy height known in the region (Figure 7a). It shows low canopy heights in the eastern half of Kruger National Park on the nutrient rich basaltic soils which are primarily grasslands with sparse trees, while canopy height tends to be greater on the nutrient poor granitic soils in the western half of the park which supports more trees and shrubs. Predicted RH98 is also higher where taller trees are known to exist such as in riparian areas (Figure 7b), along steep hillsides (Figure 7d), and especially in the western forests and tree plantations (Figure 7c). However, the height of large individual trees and small tree patches in sparse savanna areas were often underestimated (Figure 7e), and tree height in areas of grasslands and low shrubs were overestimated as 2-3 m.

**Figure 7.**
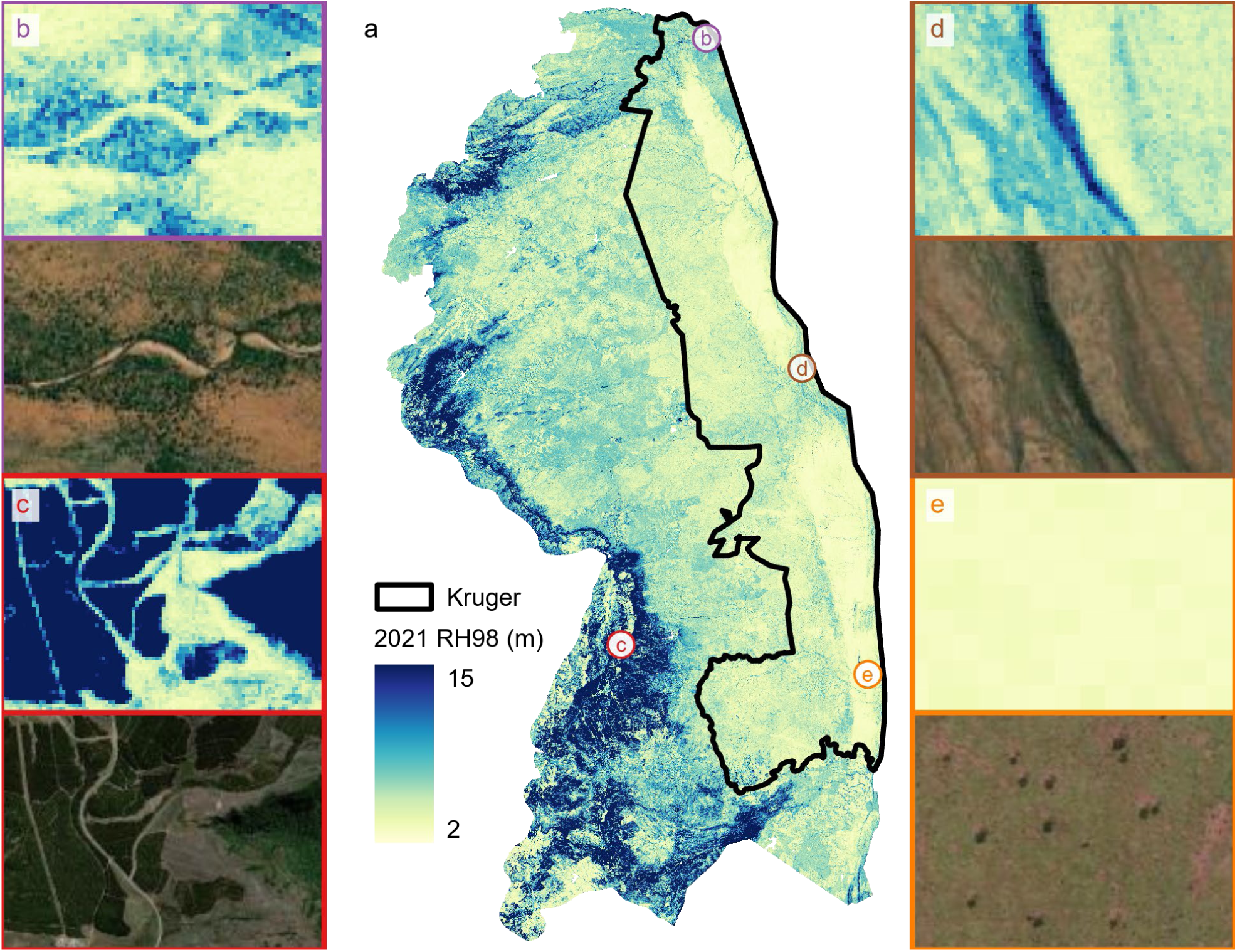
Map of predicted 2021 RH98 for the Greater Kruger region (a). Examples of the patterns of predicted canopy height and the corresponding 2021 very high resolution satellite imagery are shown for a riparian area (b), a forest and tree plantation area (c), along a steep hillside (d), and in an open area with sparse trees (e).

#### 4.2.1 Elephant Impacts in Thornybush Reserve

The map of canopy cover change in Thornybush Reserve from 2017, when eastern fences with adjacent reserves were removed, to 2021 shows a distinct decline within the property boundaries (Figure 8a). There are slight increases in cover in communal lands to the west and south where fences still exist. The decrease in canopy cover exceeded 20% in some areas (Figure 8c & d). Mean predicted canopy cover for the reserve declined from 13.0 ± 2.0% in 2017 to 8.4 ± 0.5% in 2021 (Figure 8b). Predicted RH98 and FHD showed less dramatic declines but also had relatively smaller 90% confidence intervals such that the change could still be deemed significant.

**Figure 8.**
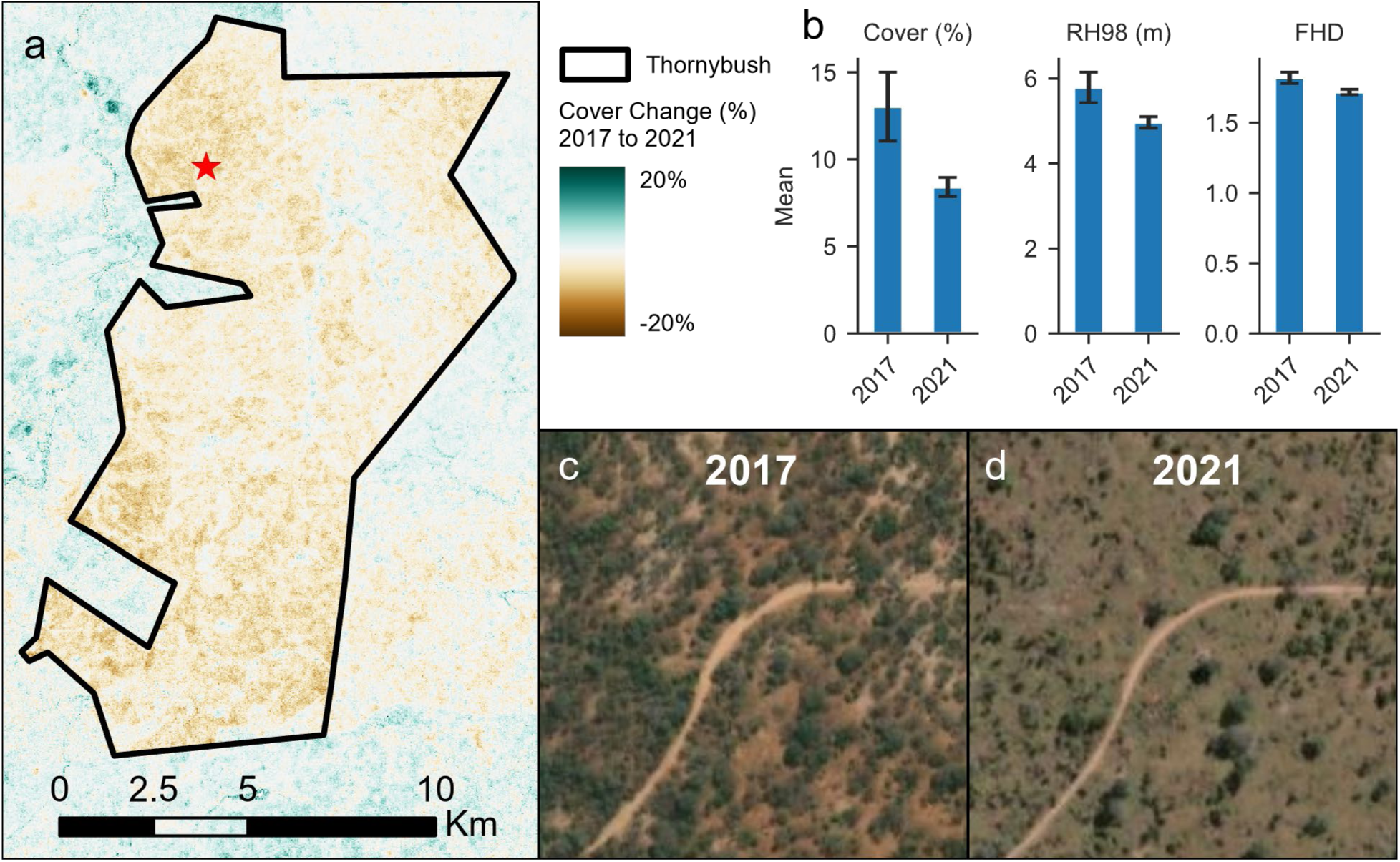
Change in predicted canopy cover in Thornybush Private Nature Reserve from 2017 to 2021 (a). Bar chart of model-based estimators of the mean cover, RH98, and FHD in 2017 and 2021 with 90% confidence interval shown as black error bars (b). Maxar satellite imagery in 2017 (c) and 2021 (d) of the area indicated by a red star in map a.

#### 4.2.2 Timber harvesting in tree plantations

Figure 9 illustrates the predicted 2017-2021 change in RH98 for a harvested patch within the region of tree plantations in southwestern portion of the Greater Kruger area. The pattern of RH98 change corresponds with the changes visible in Maxar imagery for the same time period (Figure 9c & d). Although the delineated area appears to contain some small patches of remaining trees post-harvest, estimates of the mean for all metrics in 2021 was higher than would be expected given that most of the area appears to be clear cut (Figure 9b). Mean cover also appears to be lower than what’s visible in the imagery in 2017 (Figure 9c). The confidence intervals for each metric are relatively small due to the low variance in the mapped values for each year, but this indication of uncertainty does not include error of the GEDI measurements themselves or the possibility of bias in specific areas or time periods.

**Figure 9.**
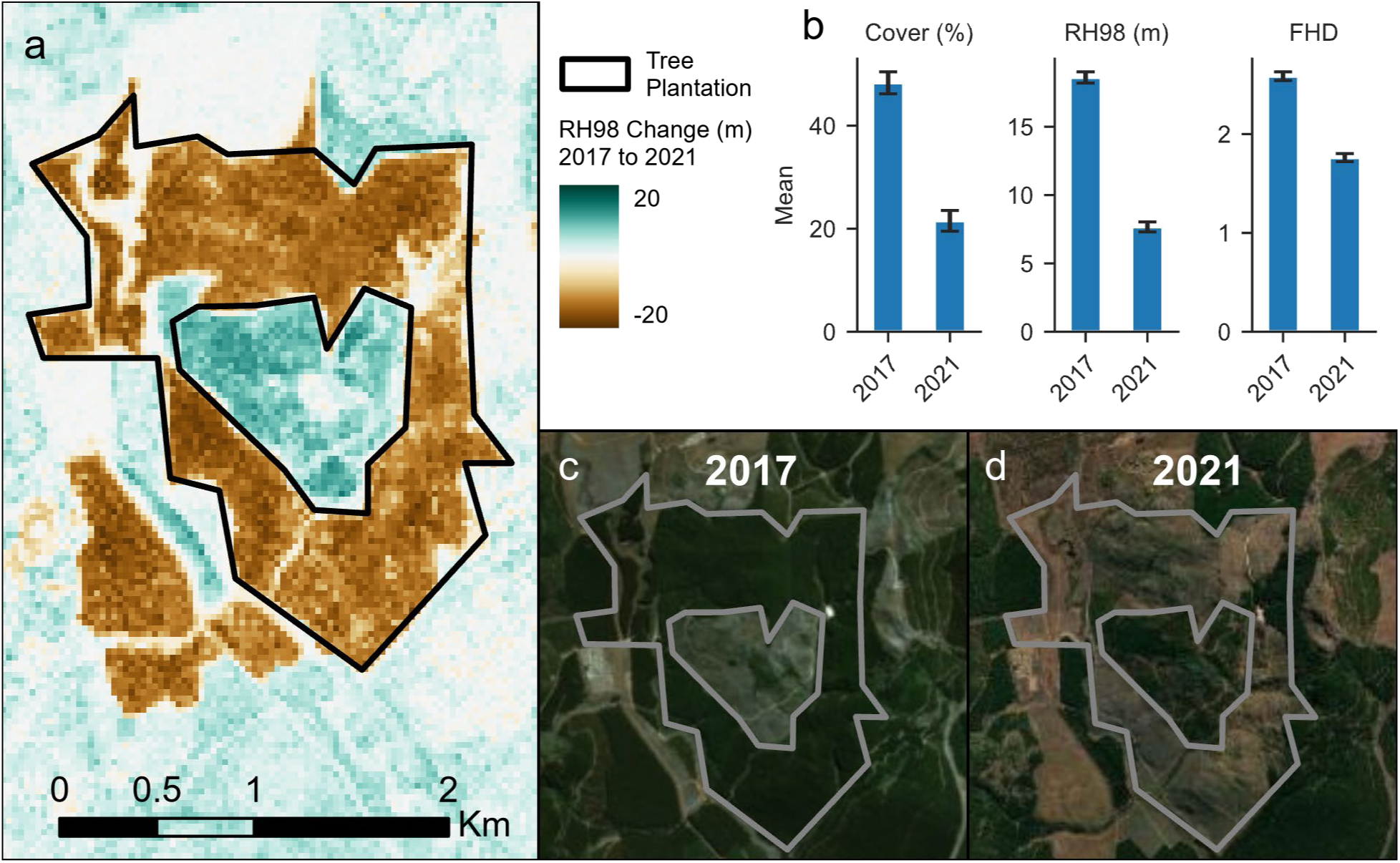
Change in predicted RH98 in a tree plantation area from 2017 to 2021 (a). Bar chart of model-based estimators of the mean cover, RH98, and FHD in 2017 and 2021 with 90% confidence intervals shown as black error bars (b). Maxar satellite imagery of the area in 2017 (c) and 2021 (d).

#### 4.2.3 Fuelwood harvesting in Bushbuckridge

The Bushbuckridge communal lands showed an increase in canopy cover from 2007 to 2021, as did the surrounding landscape (Figure 10a). The predicted increase in cover from 11.6 ± 2.0% in 2007 to 14.4 ± 2.0% in 2021 was not significant at the 90% confidence level (Figure 10b). However, there were notable changes in specific areas such as near the riparian zone that runs east to west. The pattern of increase in cover in these areas was captured by the change map and can be seen in very high-resolution satellite imagery (Figure 10c & d). Changes in RH98 and FHD were relatively less than that of canopy cover and also not significant.

**Figure 10.**
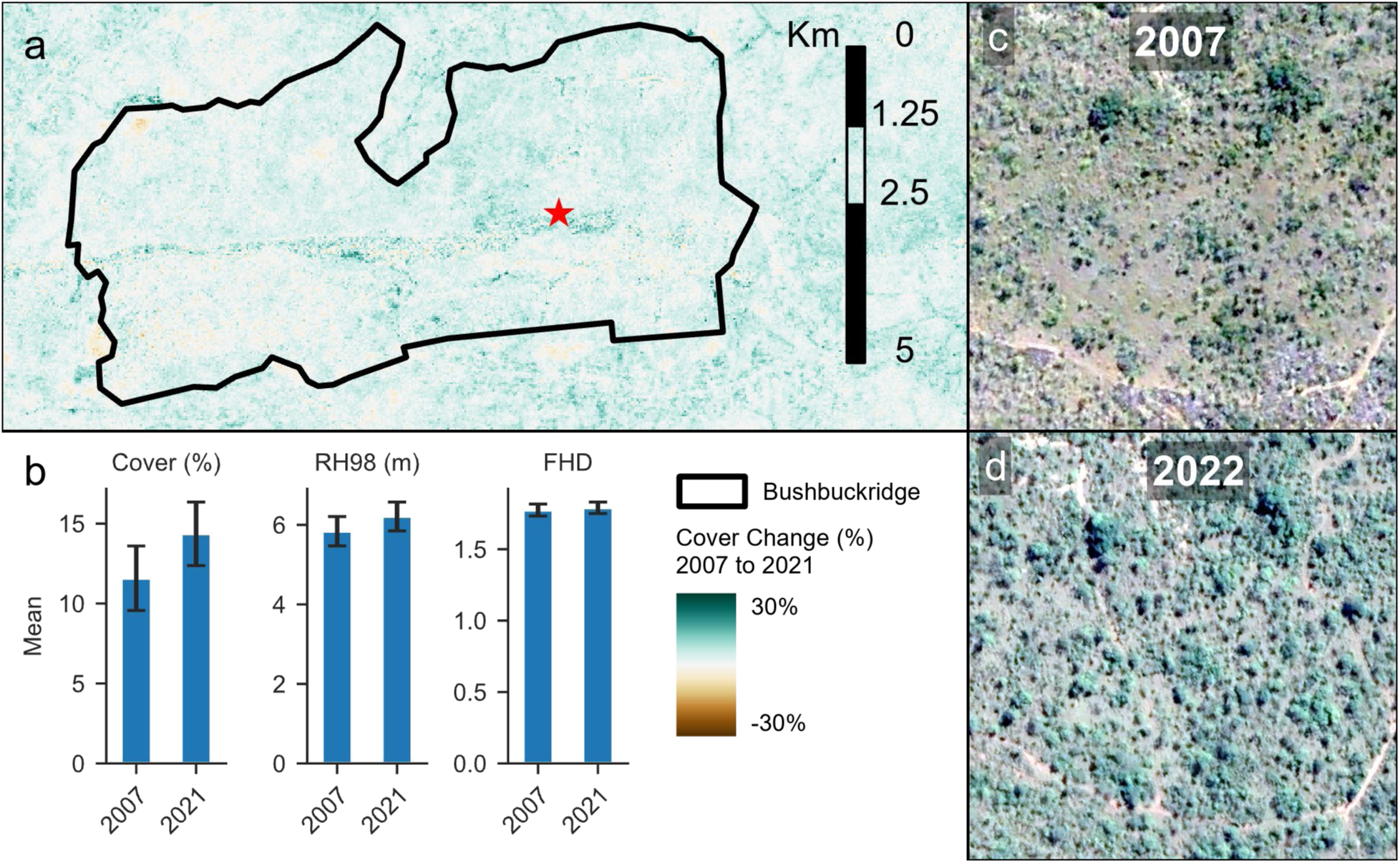
Change in predicted canopy cover in the Bushbuckridge area from 2007 to 2021 (a). Bar chart of modelbased estimators of the mean cover, RH98, and FHD in 2007 and 2021 with 90% confidence intervals shown as black error bars (b). Maxar satellite imagery in 2007 (c) and 2022 (d) of the area indicated by a red star in map a.

#### 4.2.4 Woody encroachment near Skukuza

The delineated area to the southeast of Skukuza experienced noticeable woody encroachment of shrubs between 2009 and 2024 (Figure 11c and d), as have many other parts of Greater Kruger on granitic soils. This encroachment was reflected in the map of change in canopy cover between 2007 and 2022 (Figure 11a). The mean cover increased from 7.3 ± 2.0% in 2007 to 12.5 ± 1.5% in 2022. Increases in cover appeared to be greater near drainages. There was also a significant increase in RH98 from 4.6 ± 0.4 m in 2007 to 5.9 ± 0.3 m in 2022 and in FHD from 1.66 ± 0.04 in 2007 to 1.81 ± 0.04 in 2022.

**Figure 11.**
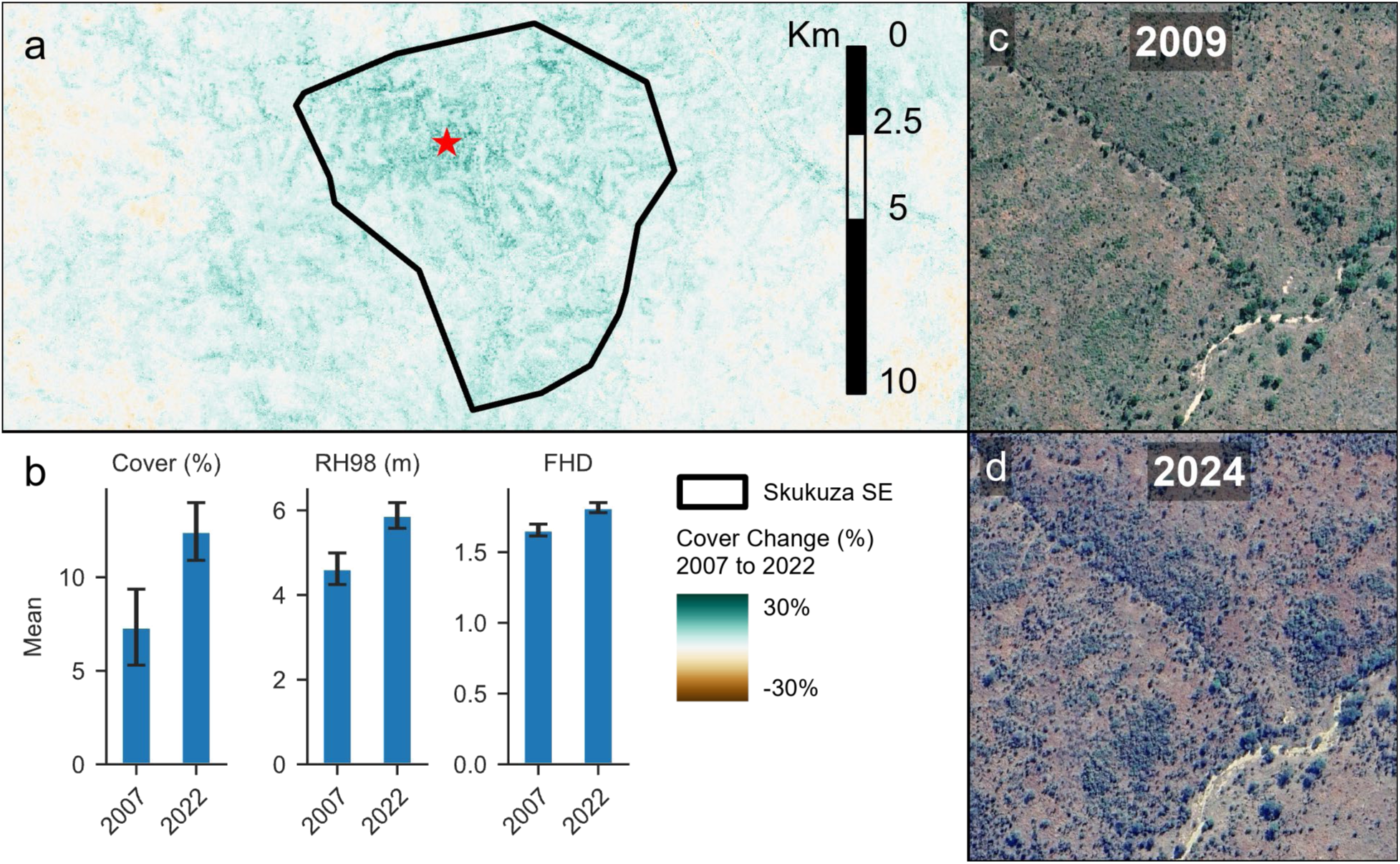
Change in predicted canopy cover in an area of woody encroachment southeast from Skukuza from 2007 to 2022 (a). Bar chart of model-based estimators of the mean cover, RH98, and FHD in 2007 and 2022 with 90% confidence intervals shown as black error bars (b). Maxar satellite imagery in 2009 (c) and 2024 (d) of the area indicated by a red star in map a.

## 5 Discussion

Our results show that fusion of moderate resolution sensors can capture canopy structure as estimated by GEDI for adequately mapping vegetation spatial patterns across a wide range of land cover and vegetation types in an African region dominated by savanna. The best performing models using a combination of moderate resolution sensors were able to explain 56%, 65%, and 50% of the variation in canopy cover, RH98, and foliage height diversity, respectively. Studies extending airborne lidar-derived woody cover with moderate resolution sensors in the Kruger area have found plot sizes of 50-100 m to yield the best trade-off in resolution and accuracy (Mathieu et al., 2013; Urbazaev et al., 2015), with the best model RMSEs ranging from 6.7% to 13.4% and R^2^ from 0.71 of 0.85 (Mathieu et al., 2013; Naidoo et al., 2016; Urban et al., 2020; Urbazaev et al., 2015; Wessels et al., 2023). Studies using higher resolutions of 15-30 m had RMSEs of 15.8% to ∼19% and R^2^ of ∼0.24 to 0.5 for woody cover, which is a closer comparison to this study given GEDI’s 25 m footprints (Mathieu et al., 2013; Urban et al., 2020; Urbazaev et al., 2015). Extension of the GEDI canopy metrics with moderate resolution sensor have obtained RMSEs of 14.6% for cover and 0.39 for FHD in North American temperate forests (Vogeler et al., 2023). We expected that the combination of GEDI’s geolocation uncertainty of ∼10 m (Li et al., 2023) and the high spatial heterogeneity of savannas would translate to lower accuracy when compared to similar scale ALS-based studies or GEDI extension in other forest types, but we instead found our model accuracies to be comparable.

Although the GEDI model results are on par with airborne lidar studies using similar scale, the reported model accuracies do not capture the additional error between GEDI observations and the ground conditions as determined from airborne lidar or field measurements. GEDI RH98 is known to have an RMSE of 1.64 m in African savannas (Li et al., 2023), but other canopy metrics have yet to be evaluated against airborne lidar in savannas specifically. The waveform return from spaceborne lidar may spread in areas of high slope, which makes ground differentiation difficult and further reduces canopy height accuracy for short-stature trees in particular (Gwenzi and Lefsky, 2014). Canopy metrics from GEDI footprints intersecting man-made structures may also misrepresent vegetation structure. Comparisons of GEDI cover to ALS-based cover measurements have found overall RMSEs of 10% to 21% and biases of ∼0% to 11% across forest types (X. Li et al., 2024; Y. Li et al., 2024), with variations in accuracy dependent on the airborne lidar cover definition and forest type among other factors. Our comparison of GEDI RH98 with a small field dataset had an accuracy similar to that of Li et al. (2023) with an RMSE of ∼2 m, and predicted RH98 had a poorer fit to field measurements (i.e., RMSE = 4.07 m) largely because of underestimation of one tall tree. Propagating errors from ground or airborne lidar measurements to GEDI to wall-to-wall maps will be particularly important for uncertainty estimation in future research in savannas because of the lower values and range of variation in canopy structure compared to most forests, which means relative error (i.e., as a percent of the prediction) could be high compared to forests.

The comparison of different predictor sets showed that fusion of multiple types of data sources improved model performance, but that including additional optical observations (i.e., HLS vs L30) or more complex temporal processing (i.e., CCDC) offered marginal benefit. While LandTrendr and CCDC have been directly compared for disturbance detection (Cohen et al., 2018), this is the first comparison that we are aware of for mapping canopy structure attributes across time. We used the LandTrendr fusion model for subsequent analyses because it provided a simpler model with fewer variables and faster computation for an increase of <1.5% RMSE across metrics compared to CCDC fusion models. However, the stability of CCDC segmentation is more robust to the addition of new observations and can yield predictions at any date (Pasquarella et al., 2022), making it potentially a better tool for frequent monitoring. Using HLS did not improve CCDC fits for each band and the HLS-based model actually performed slightly worse than when using Landsat-only for Cover and FHD, indicating that additional observations were not needed or that the true HLS dataset which features BRDF correction may be necessary. Although the use of CCDC with HLS versus Landsat-only has not previously been compared for canopy structure modeling, data reconstruction with harmonic regression improves when there are no lengthy temporal gaps and at least seven observations per year (Pasquarella et al., 2022; Zhang et al., 2021), which may be hard to obtain in regions with cloudy wet seasons. Also, use of HLS in CCDC has improved capturing of phenology, fires, and wetland dynamics in grasslands (Zhou et al., 2019). Using Landsat as the only predictor set would enable mapping of canopy structure as early as 1985 and would be only 7% less accurate on average than the best fusion models. Another interesting contrast to previous research is that the optical-only LandTrendr model explained 12% more variance than the PALSAR model, despite the higher performance of SAR-based models for mapping canopy cover in the region (Naidoo et al., 2016). The lower performance from SAR in our case may be due to using JAXA’s annual mosaics instead of careful selection of individual scenes.

Models were able to be transferred to different years with minimal impact on overall accuracy, but the compression towards the mean indicates that applying the models in areas of particularly high or low canopy cover or height likely leads to biased estimates. Temporal cross-validation had slightly lower R^2^ values and slightly higher RMSEs than out-of-bag accuracy with an absolute bias that was still close to zero for all canopy metrics. This test indicates that there is unlikely to be substantial bias or changes in error from temporal transfer to years with similar conditions (Filippelli et al., 2024). However, temporal transferability was only tested over 2018-2023 because of GEDI availability. Evaluating transferability over time spans encompassing a wider range of conditions, such as extremely wet or dry years, and checking model coverage of predictor ranges (Meyer and Pebesma, 2021) would improve user confidence in transferability. The greater source of variation in accuracy was from compression towards the mean leading to overestimation of low values and underestimation of high values as is common with many modeling approaches (Zhang and Lu, 2012). The effects of this over/under-estimation were also apparent when visually comparing maps to VHR imagery.

Maps of predicted canopy structure captured the expected broader patterns of relative height or cover without any noticeable artifacts (Figure 7). However, GEDI’s inability to estimate RH98 below 2.34 m results in maps that overestimate canopy height in open grasslands (Li et al., 2023), and because of the small sample of high values (Figure 2), predicted RH98 rarely exceeded 20 m even in riparian areas that often have trees over 25m. Predicted RH98 also underestimated canopy height in areas of very sparse trees or small patches (Figure 7e), which may be a fundamental limitation of the scale of observation (Woodcock and Strahler, 1987). Use of higher spatial resolution imagery with a frequent return, such as PlanetScope or Sentinel-2, may be able to better capture the spatial heterogeneity of tree height and cover in savannas for monitoring (Back et al., 2025).

Model-based estimators of the mean for the domains reflected the same pattern of over/under-estimation when compared to design-based estimators, though the effect was much more muted (Figure 6). The majority of model-based estimators were less than 10% different from the corresponding design-based estimator, despite having overlapping confidence intervals only ∼69-80% of the time. Since the differences between the estimators were rarely substantial, even if statistically significant, the potential bias of the model-based estimators would not negatively impact most applications assessing average canopy structure for a single year. Because bias is more likely when an area’s true mean departs from the mean of the training data (Figure 6b), the most important implication is that estimates of change may be muted when going from especially high to low values of canopy structure or vice versa. This effect could be seen in some of the case studies of change.

The direction of changes observed in the case study areas were in line with prior expectations, but the degree of change seen in maps or through model-based estimation indicate that improvements need to be made for better quantifying changes. This can be most clearly seen in the tree plantation example where mean cover decreases from 48% to 22% (Figure 9b) even though cover appears to be >70% pre-harvest and <15% post-harvest in the imagery (Figure 9c and d). The magnitude of change is less apparent in the imagery for the other case study areas. Repeat airborne lidar would be a more effective means of assessing the accuracy of change, but it is rarely available. Instead, we make comparisons to VHR imagery and existing research on the same or similar areas. Mean predicted canopy cover in Thornybush declined by 4.6% from 2017 when fences were removed to 2021 (Figure 8b), with decreases exceeding 20% in some areas. Field-based studies of other private nature reserves in the Kruger area have observed even greater elephant impacts following fence removal. The Sabi Sand Wildtuin-MalaMala complex removed fences with KNP in 1993 and witnessed a 17-fold increase in elephant populations, which resulted in a ∼60% decrease in total tree density between 1992 to 2011 (de Boer et al., 2015).

They also observed the sharpest decline in large trees, with a decrease of 261 trees ha^-1^ in the first five years after fence removal for trees >5 m tall (de Boer et al., 2015). By contrast, predicted RH98 declined by only 0.8 m from 2017 to 2021 in Thornybush (Figure 8b), even though VHR imagery would suggest a more substantial decline in the number of large trees in many areas (Figure 8c and d). Other instances of fence removal or reintroduction of elephants have similarly resulted in rapid reductions in large trees preferred by elephants, such as marula (*Sclerocarya birrea*) (Cook et al., 2017; O’Connor, 2017), and overall reductions in tree species richness, density of tall trees, and total basal area, particularly in riparian areas (O’Connor and Page, 2014). However, NDVI from AVHRR actually increased for private reserves in the Kruger area following fence removal (Linden et al., 2022), possibly due to increased greenness from woody encroachment of shrubs overriding tree losses. The relatively low overall decline in canopy height and cover we observed in Thornybush may be due in part to the underestimation of high canopy values prior to fence removal and overestimation of low canopy values following fence removal. This effect was also seen in the plantation harvest case study area (Figure 9), which was still showing a mean RH98 of 7.7 m post-harvest. Also, in very low tree density savannas, canopy height was effectively smoothed out by the moderate resolution sensors (Figure 7e), which means that residual large trees may be difficult to detect in areas where density has been reduced substantially, like in some parts of Thornybush.

There have been conflicting reports on the sustainability of fuelwood harvesting in the Bushbuckridge municipality of South Africa and its impact on vegetation structure (Shackleton et al., 2022, Table 2). Our analysis of canopy structure change in a Bushbuckridge communal rangeland showed a non-significant increase of 2.8% in cover and even less substantial relative changes in RH98 and FHD (Figure 10). These changes are in line with coppice growth from fuelwood harvesting (Twine and Holdo, 2016), but repeat lidar surveys of nearly the same area of Bushbuckridge show an even greater compensatory regrowth and bush thickening between 2008 and 2012, with a greater than 1 m increase in height for 21.1% of trees and an increase in estimated biomass of 11.34 Mg ha^-1^ (Mograbi et al., 2019, 2015). However, our maps also showed an increase in cover across much of the surrounding area (Figure 10a), suggesting the changes could alternatively be from either global causes of woody encroachment such as CO_2_ enrichment (Stevens et al., 2016), regrowth from decreases in fuelwood consumption if alternative energy sources were adopted in later years (Twine and Holdo, 2016), or simply error in the maps. Although the changes in the canopy structure maps were corroborated with VHR imagery in some areas (Figure 10c and d), either higher precision remote sensing models or a greater degree of change will be necessary to detect that a significant change has occurred in similar scenarios.

The change in canopy cover from woody encroachment was indeed detectable in other areas that had a greater magnitude of change. The area to the southeast of Skukuza had a mean increase in cover of 5.2% over 15 years, or 0.35% per year on average. Another study estimated woody encroachment in the southwestern portion of KNP using a Landsat-based model calibrated from 35 field plots and found an increase in the area of woody plants from 3.4% in 1992 to 10% in 2022, which is 0.22% per year on average (Maphanga et al., 2024). While overall woody plant cover is known to be increasing, the increase is occurring primarily in shrubs and short-stature trees like *Dichrostachys cinerea* and *Combretum apiculatum*, while larger trees like *Acacia nigrescens* and *Sclerocarya birrea* are decreasing in density (Zhou et al., 2021). We predicted an increase in canopy height in the Skukuza case study area, which could reflect the increasing area of short-stature trees. We also did not see any substantial reduction in large trees in VHR imagery for this area, as is happening primarily in areas closer to perennial water where elephants frequent.

## 6 Conclusions

The results of this study demonstrate that extension of GEDI canopy metrics through a combination of moderate resolution sensors can capture the broad patterns of canopy structure and changes in an African savanna, even when extrapolating temporally. Given that studies using airborne lidar data as a reference source yield similar accuracy when modeling at similar scales (i.e., 15-30 m), space-borne lidar presents an appealing alternative reference source for wall-to-wall mapping of canopy structure in savannas that is available globally with greater frequency. However, both the model results and maps revealed an overestimation of low values and underestimation of high values that likely resulted in biased estimators for specific areas and muted estimates of change. Improving model accuracy at the distribution’s extremes and propagating uncertainty will be crucial for advancing this approach from monitoring the direction and relative intensity of changes to reliably quantifying changes. Despite these remaining challenges, being able to map canopy structure and vegetation change over vast savanna landscapes with this approach will enable research applications that are typically confined to smaller scales or sample sizes. For example, ecologists may use these maps to examine differences in structure between vegetation types, and the maps could be included in models of wildlife occupancy and movement patterns. Time series maps of canopy structure will also allow land managers to monitor for significant changes occurring from pressures such as increasing elephant populations, illegal timber harvesting, fuelwood harvesting, and woody encroachment. Advancing regular monitoring of savanna vegetation at a high resolution will thus enable the adaptive management strategies being implemented in Kruger National Park and support policy making for the Greater Kruger region.

## 7 Acknowledgements

This study was funded by NASA Ecological Forecasting award (80NSSC21K1716) and part of the South African National Parks registered project SS610. Thank you to research and scientific support staff from SANParks, Transfrontier Africa, and the Nsasani Trust for project field support and valuable ecosystem insights.

## 8 CRediT authorship contribution statement

- **Steven Filippelli**: Conceptualization, Methodology, Software, Validation, Formal analysis, Investigation, Data Curation, Writing – Original Draft, Writing – Review & Editing, Visualization
- **Jody Vogeler**: Conceptualization, Methodology, Investigation, Resources, Writing – Review & Editing, Supervision, Project administration, Funding acquisition
- **Francisco Mauro**: Methodology, Writing – Review & Editing
- **Corli Coetsee**: Validation, Writing - Review & Editing
- **Patrick Fekety**: Methodology, Writing - Review & Editing
- **Melissa McHale**: Investigation, Supervision, Funding acquisition
- **David Bunn**: Conceptualization, Investigation, Resources, Writing – Review & Editing, Supervision, Project administration, Funding acquisition

## 9 Declaration of generative AI and AI-assisted technologies in the writing process

During the preparation of this work the author(s) used Google Gemini and OpenAI ChatGPT to assist in various aspects of writing. This included constructing a rough draft of the introduction and discussion from a detailed outline, making a rough draft of the abstract and highlights and selecting keywords based on the initial manuscript, and assistance in editing. After using this tool/service, the author(s) reviewed and edited the content as needed and take(s) full responsibility for the content of the published article.

